# The effect of pre-resection obesity on post-resection body composition after 75% small bowel resection in rats

**DOI:** 10.1101/2020.11.11.377945

**Authors:** Neesha S. Patel, Ujwal R. Yanala, Shruthishree Aravind, Roger D. Reidelberger, Jon S. Thompson, Mark A. Carlson

## Abstract

**Background:** In patients with short bowel syndrome, an elevated pre-resection Body Mass Index may be protective of post-resection body composition. We hypothesized that rats with diet-induced obesity would lose less lean body mass after undergoing massive small bowel resection compared to non-obese rats.

**Methods:** Rats (CD®IGS; age = 2 mo; N = 80) were randomly assigned to either a high-fat (obese rats) or a low-fat diet (non-obese rats), and fed ad lib for six months. Each diet group then was randomized to either underwent a 75% distal small bowel resection (massive resection) or small bowel transection with re-anastomosis (sham resection). All rats then were fed ad lib with an intermediate-fat diet (25% of total calories) for two months. Body weight and quantitative magnetic resonance-determined body composition were monitored.

**Results:** Preoperative body weight was 884 ± 95 *vs*. 741 ± 75 g, and preoperative percent body fat was 35.8 ± 3.9 *vs*. 24.9 ± 4.6%; high-fat *vs*. low fat diet, respectively (p < 0.0001); preoperative diet type had no effect on lean mass. Regarding total body weight, massive resection produced an 18% *vs*. 5% decrease in high-fat vs. low-fat rats respectively, while sham resection produced a 2% decrease *vs*. a 7% increase, respectively (p < 0.0001, preoperative *vs*. necropsy data). Sham resection had no effect on lean mass; after massive resection, both high-fat and low-fat rats lost lean mass, but these changes were not different between the latter two rat groups.

**Conclusion:** The high-fat diet and low-fat diet induced obesity and marginal obesity, respectively. The massive resection produced greater weight loss in high-fat rats compared to low-fat rats. The type of dietary preconditioning had no effect on lean mass loss after massive resection. A protective effect of pre-existing obesity on lean mass after massive intestinal resection was not demonstrated.

## Introduction

The prevalence of short bowel syndrome (SBS) in United States is in the range of 30-45 per million,^1,2^ but SBS may be under-reported due to lack of a national registry or a database and variation in definition. The incidence often is estimated from home parenteral nutrition registries.^2^ A Swedish SBS registry estimated an SBS prevalence of 44 per million in that country.^3^ Approximately 15% of adults who undergo massive intestinal resection will suffer from SBS.^4,5^ A process of structural and functional adaptation occurs in SBS in which the intestinal remnant hypertrophies and dilates, thereby increasing the absorptive capacity of the intestines.^6^ In addition, a distinct process called *adaptive hyperphagia* has been described in patients with SBS, wherein the dietary intake is markedly increased as a compensatory mechanism for reduced absorptive capacity.^7^

Clinical studies have suggested that the pre-resection body mass index (BMI) influences post-resection BMI in patients with SBS.^8,9^ Obesity may affect the intestinal adaptive response, energy expenditure, body composition and adaptive hyperphagia after intestinal resection.^9^ In order to determine whether pre-existing obesity would be protective of body weight and other nutritional endpoints in experimental SBS, we previously^10^ performed a study on the effect of a 50% small intestinal resection on nutritional endpoints in rats with and without diet-induced obesity. The obese rats may have retained more lean body mass after resection that non-obese rats, but the difference was small and not statistically significant.^10^ Herein we report a repeat of the above rat study, but this time using a greater intestinal resection (75% of the small bowel) and a longer postoperative follow-up period (two months, as opposed to the previous one-month period). The rationale for this repeat study was to follow up on the inconclusive results of the previous study,^10^ using a more severe model of SBS to determine whether rats with diet-induced obesity would have greater preservation of lean mass after massive resection than non-obese rats.

## Materials and Methods

### Animal Welfare

This study was performed with approval from the Institutional Animal Care and Use Committees of the VA Nebraska-Western Iowa Health Care System (protocol number 00992) and of the University of Nebraska Medical Center (protocol number 14-082-09-ET), and in accordance with the recommendations in the NIH *Guide for the Care and Use of Laboratory Animals* (8th ed.).^11^ Procedures were performed and subjects housed in animal facilities approved by the Association for Assessment and Accreditation of Laboratory Animal Care International (AAALAC; www.aaalac.org). Animals were housed in pairs, except for a period of single housing for 7 days after surgery. Surgical procedures were performed under isoflurane anesthesia, and all efforts were made to minimize pain and distress. Buprenorphine SR (0.1 mg/kg injection) was given as needed or pain. Any subject in distress, not eating, not moving, or with obvious wound infection was euthanized. Euthanasia was performed in accordance with the AVMA Guidelines.^12^ The ARRIVE 2.0 Guidelines (Animal Research: Reporting of *In Vivo* Experiments^13,14^) Checklist is shown in Fig. S1.

### Animal Subjects and Determination of Subject Numbers

CD® IGS (International Genetic Standard) rats (all males; age 3 mo upon arrival) were purchased from Charles River Laboratories International, Inc. (The minimum number of rats (N = 12) utilized for each experimental group (four groups total; see Fig. 1) was determined with a statistical power analysis using Δ/σ (= Cohen’s *d*, in which Δ is the desired difference in means as set by the observer, and σ is the estimated standard deviation) = 1.50, false positive rate (α) = 0.05, false negative rate (β) = 0.2, and power (1 – β) = 0.8. With N = 12 rat/group and α and β set as above, a 20% difference among means could be detected with a standard deviation equal to 13% of the mean. The primary endpoint in this study was lean body mass. The actual number of subjects per group at the beginning of the dietary induction phase in Fig. 1 was increased to 15. The justification for this increased subject number was in case of subject loss secondary to illness or perioperative mortality.

**Fig. 1.**
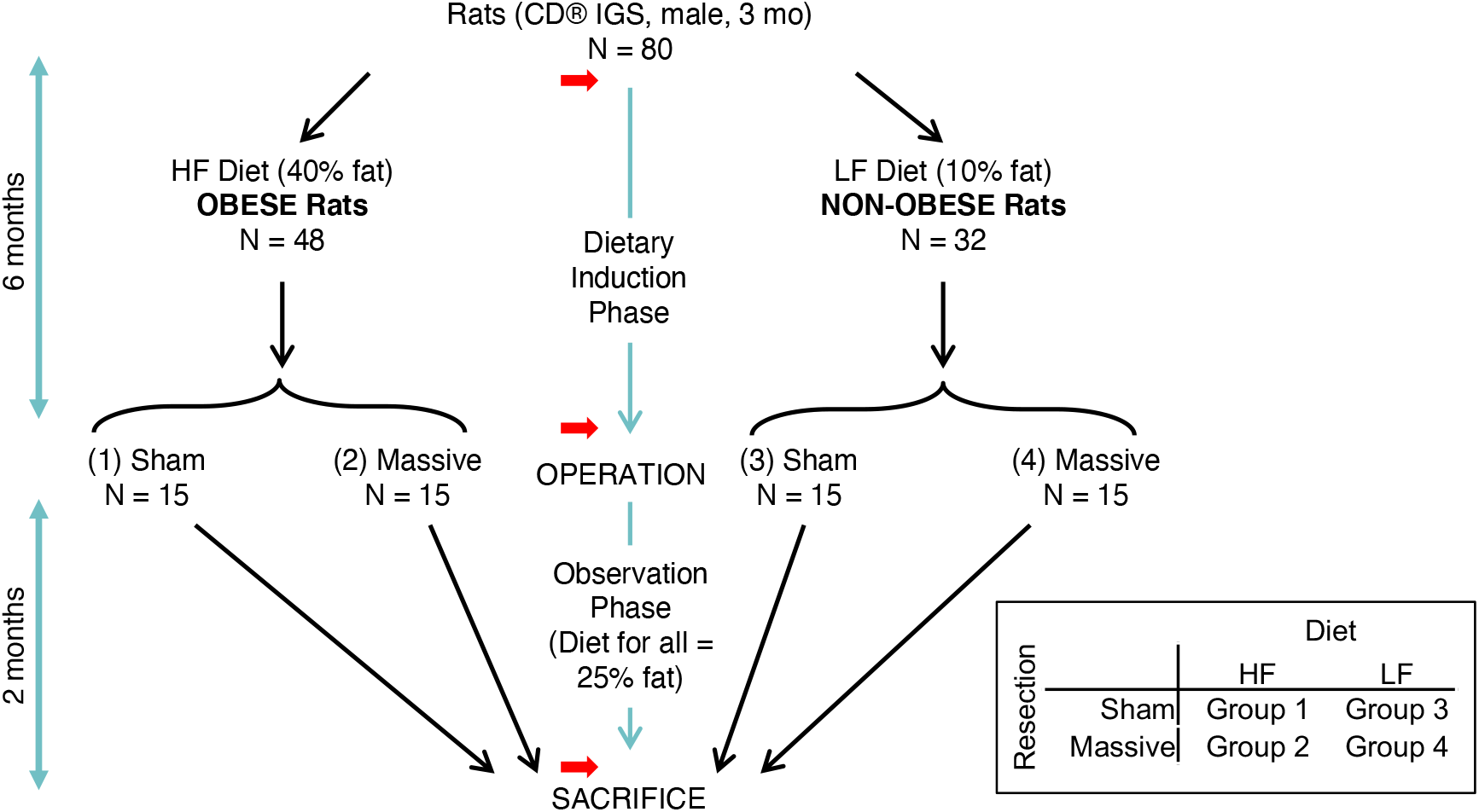
Experimental Design. HF = High-Fat; LF = Low-Fat; Sham = sham resection; Massive = 75% distal small intestinal resection. Red arrow = body composition measurement. Inset: 2 x 2 matrix of experimental groups.

### Experimental Design Overview

The experimental design (Fig. 1) was adapted from our previous study.^10^ Rats were assigned to either a High-Fat (HF) or a Low-Fat (LF) diet for 120 days, and then underwent either a 75% distal enterectomy (“massive resection”) or a transection of the distal ileum with anastomosis (“sham resection”). This design yielded a 2 x 2 matrix of experimental groups (Fig. 1). Postoperatively all animals were maintained on an Intermediate-Fat (IF) diet for 60 days, and then euthanized. Body composition was measured prior to dietary induction, prior to operation, and prior to sacrifice (Fig. 1).

### Dietary Induction Phase

After a one week acclimatization period on standard rat chow, body composition measurement was obtained and rats (N = 80) were randomly assigned to either the HF diet (45% of total cal from fat; OpenSource Diets® product no. D12451; researchdiets.com; see Table 1 and Fig. S2; N = 48 rats) or the LF diet (10% of total cal from fat; OpenSource Diets™ product no. D12450B; researchdiets.com; see Table 1 and Fig. S3; N = 32 rats), using an online randomizer (www.randomizer.org). The justification for the greater number of rats assigned to the HF diet was the anticipation that some subjects on this diet would not become obese.^10^ At the time of randomization to the operative procedure, there would need to be 30 rats in each diet group (see Fig. 1); so, the starting number of rats on the HF diet needed to be well over 30. Rats were caged in pairs, and fed their assigned diet *ad libitum* for 6 months, which was defined as the dietary induction phase (Fig. 1).

**Table 1.** Dietary formulations.

| Quantity | High-Fat Diet<sup>*</sup> | Low-Fat Diet<sup>†</sup> | Intermediate-Fat Diet<sup>#</sup> |
| --- | --- | --- | --- |
| Energy density (kcal/g) | 4.73 | 3.85 | 4.73 |
| Protein kcal (%) | 20 | 20 | 24 |
| Carbohydrate kcal (%) | 35 | 70 | 41 |
| Fat kcal (%) | 45 | 10 | 25 |

High-fat diet<sup>\*</sup> = D12451; low fat diet<sup>†</sup> = D12450B (both from OpenSource Diets<sup>®</sup>); intermediate fat diet<sup>#</sup> = LabDiet 5001 Rodent Diet (PMI Nutrition<sup>®</sup> Inc.). See Supplementary Information for more diet details.

### Surgical Procedure

Surgical procedures were based on a prior description.^10^ After the dietary induction phase, body composition measurement was obtained, and within each group the 30 rats with the highest body weight were then randomly assigned to either massive or sham resection (Fig. 1), using the above online randomizer. Rats were fasted for 24 h prior to operation, but with free access to water. Anesthetic induction was obtained with 5% isoflurane via a nose cone and maintained with 2% isoflurane throughout the surgical procedure.^10^ After ensuring adequate anesthesia, subcutaneous cefovecin (20mg; a long acting cephalosporin) was administered, the ventral hair was clipped, the abdomen was scrubbed with antiseptic solution (Povidone Iodine Topical Solution, 10% USP, Allegiance, Cardinal Health), and the surgical field was isolated with sterile disposable drapes. A 6-cm midline laparotomy from the xiphoid process to urethral meatus was performed. The small intestine was exteriorized, and the entire length (from the pylorus to the cecum) was measured using a cotton umbilical tape. Intestinal surfaces were kept moist during the procedure with saline-soaked gauze.

Rats assigned to massive resection (Groups 2 and 4) underwent a 75% distal enterectomy (Fig. S4-S6). The proximal transection line was in the mid jejunum, at the junction of the proximal one-quarter and distal three-quarters of the small intestinal length (Fig. S4). The distal transection line was in the distal ileum, 1 cm proximal to cecum (i.e., the ileocecal intersection was preserved). Mesenteric vessels were controlled using 4-0 silk (Fig. S5). An end-to-end jejunoileal anastomosis was performed using a single layer of running sutures with 6-0 silk (Ethicon Perma-Hand® Silk, catalog no. K889H; see Fig. S6 for technical details). Rats assigned to sham resection (Groups 1 and 3) underwent a transection of the mid jejunum at the junction of the proximal one-quarter and distal three-quarters of the small intestinal length, without resection of any bowel (Fig. S4). The transected jejunum simply was re-connected with an end-to-end anastomosis as above.

The musculoaponeurotic layer of the abdominal wall was closed with a single running 3-0 polyester suture (Ethibond Excel®, catalog no. X558H). Just prior to tying the polyester suture, 10 mL of saline was injected into the peritoneal cavity for postoperative hydration. The skin then was closed with staples (Autoclip® 9 mm stainless steel clips; Mikron Precision, Inc., catalog no. 427631). A one-time application of topical antibiotic ointment (Fura-Zone, 0.2% nitrofurazone; Squire Laboratories, Inc.) was spread along the skin staple line. An injection of long-acting buprenorphine (0.1 mg subcutaneously) was given prior to recovery from anesthesia.

### Postoperative Observation Phase

After the surgical procedure, all rats were fed *ad libitum* with an Intermediate-Fat (IF) diet (25% of total cal from fat; OpenSource Diets® product no. D14041301; researchdiets.com; see Table 1 and Fig. S7). Body composition measurement was obtained at the end of the 60-day observation phase (Fig. 1). Euthanasia then was performed under deep isoflurane anesthesia with a laparotomy, bilateral diaphragm incision, and exsanguination by cardiotomy. The small intestine from the stomach to the cecum was resected *en bloc*, grossly inspected, measured for length, and photographed.

### Body Composition Measurement

Body composition was determined noninvasively with quantitative magnetic resonance (QMR)^10^ prior to the dietary induction phase, prior to the surgical procedure, and then prior to euthanasia, using an EchoMRI™ 700 Whole Body Composition Analyzer (Echo Medical Systems; echomri.com).

### Statistical Analysis

Data was analyzed using IBM SPSS Statistics for Windows, Version 22.0. Armonk, NY: IBM Corp. All results were expressed as mean ± SD. ANOVA and t-testing were used to groups of data. Significance was defined as p < 0.05.

## Results

At the initiation of the study, the weight and body fat of the rats assigned to HFD (N = 48) and LFD (N = 32) just prior to the dietary induction phase were not different (Table 2 and Fig. S8). After the 6-month dietary induction phase, HFD and LFD rats were randomized to four intestinal resection groups (1 = HFD sham resection, 2 = HFD massive resection, 3 = LFD sham resection, 4 = LFD massive resection; see Fig. S9A), following the design shown in Fig.1. After completion of the 6-month dietary induction phase, the average weight, body fat, and percent body fat of HFD rats was greater than the LFD diet rats (Table 2); the former group gained an average of ∼500 g of weight and ∼277 g of fat, while the latter had respective gains of ∼352 g and ∼144 g. The lean mass of the HFD vs. LFD rats was not different at the end of the 6-month dietary induction phase (Table 3).

**Table 2.** Body weight and fat prior to dietary induction.

| Quantity | High-Fat Diet<br>(N = 48) | Low-Fat Diet<br>(N = 32) | p-value* |
| --- | --- | --- | --- |
| Body weight (g) | 385 ± 16 | 389 ± 20 | 0.310 |
| Body fat (g) | 41.2 ± 4.7 | 42.0 ± 5.7 | 0.474 |

Values obtained just prior to 6-month dietary induction phase. Data = mean ± SD; \*unpaired t-test.

**Table 3.** Preoperative body composition, high-fat *vs*. low-fat diet.

| Quantity | High-Fat Diet<br>(N = 30) | Low-Fat Diet<br>(N = 30) | p-value* |
| --- | --- | --- | --- |
| Body weight (g) | 884 ± 95 | 741 ± 75 | <0.0001 |
| Body fat (g) | 318 ± 59 | 186 ± 45 | <0.0001 |
| Body fat (%) | 35.8 ± 3.9 | 24.9 ± 4.6 | <0.0001 |
| Lean weight (g) | 566 ± 54 | 555 ± 53 | 0.416 |
| Lean weight (%) | 64.2 ± 3.9 | 75.1 ± 4.6 | <0.0001 |

Values obtained after 6-month dietary induction phase, immediately prior to resection. Data = mean ± SD; \*unpaired t-test.

Four rats died after the surgical procedure secondary to anastomotic leakage or intraabdominal sepsis (6.7% operative mortality), leaving ≥13 rats in each group for analysis (Table 4). The effect of diet and resection on total body weight (TBW) is shown in Table 5. Comparing all four treatment groups at single time points, there was no statistical difference in TBW among the four treatment groups neither prior to resection nor at two months after resection (i.e., at necropsy). Performing paired comparisons within each group across time (from the preoperative to the necropsy time point), there was a large (18%) decrease in TBW in group 2 (HFD massive resection), with a marginal (2%) decrease in TBW in group 1 (HFD sham resection). There was a modest (5%) decrease in TBW in group 4 (LFD massive resection), with a modest (7%) increase in TBW in group 3 (LFD sham resection). The decrease in TBW experienced by group 2 (HFD massive resection) was approximately four-fold greater than in group 4 (LFD massive resection). In the sham resection groups, there was a similar four-fold difference in TBW change between groups 1 and 3, favoring the latter (LFD sham resection). A flow diagram summarizing the fate of all 80 rats in this report is shown in Fig. S10.

**Table 4.** Operative mortality.

| Group | Operated (N) | Died (N) | Left for analysis (N) |
| --- | --- | --- | --- |
| 1 (HFD, Sham) | 15 | 1 | 14 |
| 2 (HFD, Massive) | 15 | 1 | 14 |
| 3 (LFD, Sham) | 15 | 2 | 13 |
| 4 (LFD, Massive) | 15 | 0 | 15 |
| Sum | 60 | 4 | 56 |

HFD = High-Fat Diet; LFD = Low-Fat Diet; Sham = sham resection; Massive = 75% distal small intestinal resection.

**Table 5.** Effect of diet and resection on total body weight.

|  | Total Body Weight |  |  |  |  |  |
| --- | --- | --- | --- | --- | --- | --- |
| | Pre-Diet (g) | Preop (g) | Necrop (g) | $\Delta$ (Change),<br>Necrop – Preop<br>(g) | $\Delta\%$<br>(% of Preop) | Paired t-test<br>(Preop vs. Necrop) |
| Mean, Group 1<br>(HFD Sham) | 379.4 ± 18.6 | 877.0 ± 112.3 | 858.5 ± 104.7 | (18.5) ± 36.0 | (1.9) ± 4.0 | 0.038 |
| Mean, Group 2<br>(HFD Massive) | 388.5 ± 17.3 | 880.7 ± 89.0 | 717.6 ± 97.4 | (163.1) ± 80.1 | (18.4) ± 8.4 | <0.0001 |
| Mean, Group 3<br>(LFD Sham) | 389.4 ± 23.1 | 738.4 ± 75.2 | 789.0 ± 87.8 | 50.7 ± 27.6 | 6.8 ± 3.7 | <0.0001 |
| Mean, Group 4<br>(LFD Massive) | 383.2 ± 14.5 | 718.6 ± 63.2 | 681.7 ± 69.5 | (36.9) ± 29.5 | (5.2) ± 4.1 | 0.00013 |
| ANOVA,<br>Groups 1 – 4 | 0.45 | 0.27 | 0.22 | <0.0001 | <0.0001 |  |
| Unpaired t-test |  |  |  |  |  |  |
| 1 v. 2 |  |  |  | <0.0001 | <0.0001 |  |
| 3 v. 4 |  |  |  | <0.0001 | <0.0001 |  |
| 1 v. 3 |  |  |  | <0.0001 | <0.0001 |  |
| 2 v. 4 |  |  |  | <0.0001 | <0.0001 |  |

Values shown are mean ± standard deviation. Red values in parentheses (xx.x) represent a negative change (decrease). Pre-Diet = prior to the 6-mo dietary induction period; Preop = Preoperative (after dietary induction, immediately prior to resection); Necropsy = immediately prior to necropsy (two months after resection); HFD = High-Fat Diet; LFD = Low-Fat Diet; Massive = Massive 75% distal small intestinal resection; Sham = Sham resection.

The effect of diet and resection on body fat (BF) is shown in Table 6. Comparing all four treatment groups at a single time point, group 1 and 2 (HFD rats) had greater BF than groups 1 and 3 (LFD rats) at the preoperative time point. Performing paired comparisons within each group across time (from the preoperative to the necropsy time point), there was a large (45%) decrease in BF in group 2 (HFD massive resection), with no significant decrease in BF in group 1 (HFD sham resection). There was a smaller (15%) decrease in BF in group 4 (LFD massive resection), with a large (27%) increase in BF in group 3 (LFD sham resection). The decrease in BF experienced by group 2 (HFD massive resection) was approximately five-fold greater than in group 4 (LFD massive resection). In the sham resection groups, there was a similar five-fold difference in BF change between groups 1 and 3, favoring the latter (LFD sham resection).

**Table 6.** Effect of diet and resection on body fat.

|  | Body Fat Mass |  |  |  |  | Body Fat Mass as percent of TBW |  |  |
| --- | --- | --- | --- | --- | --- | --- | --- | --- |
| | Preop (g) | Necrop (g) | $\Delta$ (Change),<br>Necrop – Preop<br>(g) | $\Delta\%$<br>(% of Preop) | Paired t-test<br>(Preop vs. Necrop) | Preop<br>%Body Fat | Necrop<br>%Body Fat | Paired t-test<br>(Preop v. Necrop) |
| Mean, Group 1<br>(HFD Sham) | 315.8 ± 69.3 | 306.7 ± 62.6 | (9.1) ± 31.2 | (1.9) ± 9.4 | 0.15 | 35.7 ± 4.8 | 35.5 ± 4.5 | 0.39 |
| Mean, Group 2<br>(HFD Massive) | 315.6 ± 50.4 | 177.1 ± 61.4 | (138.5) ± 38.2 | (44.9) ± 13.7 | <0.0001 | 35.7 ± 3.4 | 24.2 ± 6.0 | <0.0001 |
| Mean, Group 3<br>(LFD Sham) | 189.8 ± 47.9 | 239.9 ± 56.1 | 50.1 ± 19.0 | 26.9 ± 10.3 | <0.0001 | 25.5 ± 4.7 | 30.1 ± 5.1 | <0.0001 |
| Mean, Group 4<br>(LFD Massive) | 180.8 ± 43.2 | 155.6 ± 48.4 | (25.2) ± 26.5 | (14.7) ± 15.4 | 0.0012 | 25.0 ± 4.9 | 22.6 ± 5.8 | 0.0066 |
| ANOVA,<br>Groups 1 – 4 | <0.0001 | <0.0001 | <0.0001 | <0.0001 |  | <0.0001 | <0.0001 |  |
| Unpaired t-test |  |  |  |  |  |  |  |  |
| 1 v. 2 | 0.50 | <0.0001 | <0.0001 | <0.0001 |  | 0.5 | <0.0001 |  |
| 3 v. 4 | 0.30 | 0.00011 | <0.0001 | <0.0001 |  | 0.41 | 0.00066 |  |
| 1 v. 3 | <0.0001 | 0.0037 | <0.0001 | <0.0001 |  | <0.0001 | 0.0036 |  |
| 2 v. 4 | <0.0001 | 0.15 | <0.0001 | <0.0001 |  | <0.0001 | 0.24 |  |

Values shown are mean ± standard deviation. Red values in parentheses (xx.x) represent a negative change (decrease). Preop = Preoperative (after dietary induction, immediately prior to resection); Necropsy = immediately prior to necropsy (two months after resection); HFD = High-Fat Diet; LFD = Low-Fat Diet; Massive = Massive 75% distal small intestinal resection; Sham = Sham resection; TBW = Total Body Weight.

The effect of diet and resection on lean mass (LM) is shown in Table 7. Comparing all four treatment groups at single time points, there was no statistical difference in LM among the four treatment groups neither prior to resection nor at two months after resection (i.e., at necropsy). Performing paired comparisons within each group across time (from the preoperative to the necropsy time point), there was a modest (4%) decrease in LM in group 2 (HFD massive resection), with no statistical change in LM in group 1 (HFD sham resection). There also was a modest (3%) decrease in LM in group 4 (LFD massive resection), with no statistical change in LM in group 3 (LFD sham resection). Of note, the change in LM from the preoperative to the necropsy time point between group 2 (HFD massive resection) vs. group 4 (LF massive resection) was not statistically different.

**Table 7.** Effect of diet and resection on lean mass.

|  | Body Lean Mass |  |  |  |  | Lean Mass as percent of TBW |  |  |
| --- | --- | --- | --- | --- | --- | --- | --- | --- |
| | Preop (g) | Necrop (g) | $\Delta$ (Change),<br>Necrop – Preop<br>(g) | $\Delta\%$<br>(% of Preop) | Paired t-test<br>(Preop vs. Necrop) | Preop<br>%Lean Mass | Necrop<br>%Lean Mass | Paired t-test<br>(Preop v. Necrop) |
| Mean, Group 1<br>(HFD Sham) | 482.8 ± 47.3 | 480.6 ± 48.9 | (2.1) ± 18.0 | (0.4) ± 3.8 | 0.33 | 55.4 ± 4.0 | 56.3 ± 4.5 | 0.080 |
| Mean, Group 2<br>(HFD Massive) | 493.1 ± 35.2 | 472.7 ± 41.3 | (20.5) ± 14.5 | (4.2) ± 3.0 | <0.0001 | 56.3 ± 5.0 | 66.5 ± 6.2 | <0.0001 |
| Mean, Group 3<br>(LFD Sham) | 479.8 ± 41.7 | 480.8 ± 44.8 | 1.0 ± 11.3 | 0.2 ± 2.3 | 0.37 | 65.2 ± 4.1 | 61.2 ± 4.7 | <0.0001 |
| Mean, Group 4<br>(LFD Massive) | 482.0 ± 42.8 | 466.0 ± 47.6 | (16.0) ± 12.6 | (3.4) ± 2.8 | 0.00011 | 67.2 ± 4.2 | 68.5 ± 5.5 | 0.075 |
| ANOVA,<br>Groups 1 – 4 | 0.84 | 0.79 | 0.0004 | 0.0004 |  | <0.0001 | <0.0001 |  |
| Unpaired t-test |  |  |  |  |  |  |  |  |
| 1 v. 2 |  |  | 0.0032 | 0.0032 |  | 0.29 | <0.0001 |  |
| 3 v. 4 |  |  | 0.00047 | 0.00053 |  | 0.11 | 0.00046 |  |
| 1 v. 3 |  |  | 0.30 | 0.32 |  | <0.0001 | 0.0046 |  |
| 2 v. 4 |  |  | 0.19 | 0.22 |  | <0.0001 | 0.18 |  |

Values shown are mean ± standard deviation. Red values in parentheses (xx.x) represent a negative change (decrease). Preop = Preoperative (after dietary induction, immediately prior to resection); Necropsy = immediately prior to necropsy (two months after resection); HFD = High-Fat Diet; LFD = Low-Fat Diet; Massive = Massive 75% distal small intestinal resection; Sham = Sham resection; TBW = Total Body Weight.

## Discussion

This study was a second test of the hypothesis that pre-existing obesity would preserve lean mass in a rat model of massive intestinal resection (i.e., surgically-induced SBS). The first test of this hypothesis was published in 2015,^10^ and utilized a 50% small intestinal resection with a postoperative observation period of one month. This prior study did not convincingly demonstrate a protective effect of obesity; interestingly, no differential effect of 50% enterectomy vs. sham resection on body composition was demonstrable in this 2015 study.^10^ So in the second test of the above hypothesis (i.e., the present report), the size of the intestinal resection and the length of the postoperative observation period were increased to 75% and two months, respectively. The expectation for these changes in study design was that any protective effect of obesity on lean mass in this revised model would be more pronounced and possibly significant.

The data in the present report did not prove the hypothesis either; pre-existing obesity in a rat model of 75% distal small intestinal resection was not protective of lean mass two months after resection. Secondary conclusions that could be drawn from the data include: (1) the high-fat diet induced obesity (36% body fat); (2) the low-fat diet resulted in marginal obesity (25% body fat); (3) massive resection produced substantially greater weight loss than sham resection for rats on either diet; and (4) rat pre-conditioned with the high-fat diet lost more weight and body fat after massive resection compared to the low-fat diet rats. Of these secondary conclusions, numbers 1, 3, and 4 were expected outcomes, and support the validity of the study. In particular, the present study (which utilized a 75% enterectomy) demonstrated a large effect of resection on body composition in comparison to sham resection; the prior 2015 study (which utilized a 50% enterectomy), was unable to demonstrate such a differential effect.^10^

Regarding secondary conclusion number 2, the low-fat diet rats in the present study had greater percent body fat (25%) compared to the 18% body fat of similarly-preconditioned rats from the 2015 study (same rat strain).^10^ It is not clear why the low-fat diet rats in the present study attained this relatively high preoperative percent body fat, or whether these rats were “lean enough” to qualify as a “non-obese” comparator group for the rats on the high-fat diet. Experimental obesity in rats has been defined as a percent body fat of 20-25%,^15-17^ so the “non-obese” rats in the present study were on the margin of this definition. It is interesting to note, however, that low-fat diet rats in the present study actually gained weight and body fat after sham resection, while the high-fat diet rats did not; this result suggests a difference in the underlying response to stress between rats on these two diets. The relationship of this observation to the central hypothesis, which focused on lean mass, is not obvious.

The data from the present study do not support the “obesity paradox”, in which better outcomes have been hypothesized for obese patients compared to non-obese patients suffering from a variety of diseases or conditions.^18-25^ From this perspective, the present report could be classified as a “negative” study. Possible causes for the finding of no support for the obesity paradox include: (1) the non-obese rats were not “lean enough” to be a comparator group (see above discussion); (2) the rats were too old (about one year) at the study’s conclusion to mount a vigorous response to surgical stress; (3) the rat may not adequately reproduce human physiology in this area, i.e., the rat may have been inadequate model; (4) the present study did not utilize physical fitness as a variable, which may be a relevant parameter in the obesity paradox;^25^ or (5) the wrong endpoint (lean mass) was chosen for this study. Alternatively, this study may be clinically accurate in that the obesity paradox may not apply to massive intestinal resection and/or SBS.

A drawback of this study is that all of the rat subjects were male. The NIH has mandated that subject sex should be considered as a biological variable, and that grant applicants should balance male and female subjects in their preclinical investigations.^26^ Of note, the study was designed and executed prior to the NIH’s announcement of its intent to focus on sex as a biological variable. Regardless of that, the present study cannot discriminate any influence of subject sex on outcome, because there were no female subjects in this study.

Regarding the selection of an appropriate primary endpoint, in clinical cases of massive intestinal resection or SBS, relevant patient outcomes include survival, operative morbidity, nutritional parameters (including lean mass), long-term dependence on parenteral nutrition, and quality of life indicators. We selected lean mass as the best and most practical endpoint that was available to us for the rat model; this endpoint has been commonly employed in other studies on nutrition and metabolism in rats.^27-31^ In addition to the evaluation of lean mass as the primary endpoint, there are secondary endpoints that could have been evaluated in the present study, including food intake, intestinal adaptation, immunohistochemical staining, serum peptide levels, and other nutrition-relevant serum tests; of note, most of these secondary endpoints were examined in the 2015 study.^10^ We elected not to perform these analyses in the current study, because: (1) the relevance of these secondary endpoint data would not be clear, as the study was negative with respect to the primary endpoint; and (2) we did not want to expend resources in the pursuit of costly secondary endpoint data which would have doubtful relevance.

In summary, this was a report of a second study in which the effect of pre-resection obesity on preservation of lean body mass after massive intestinal resection was tested in rats. In this second study, a longer postoperative observation period and a greater intestinal resection was utilized compared to the first study. Neither study was able to demonstrate a protective effect of obesity on lean mass after massive intestinal resection.

## Acknowledgements

This study is the result of work supported in part with resources and the use of facilities at the Omaha VA Medical Center (Nebraska-Western Iowa Health Care System). The authors would like to acknowledge the technical assistance of Gerri Siford and Chris Hansen. This work was funded through the Department of Surgery at the University of Nebraska Medical Center.

