## Supplementary material for "The effect of pre-resection obesity on post-resection body composition after 75% small bowel resection in rats": Fig. S1-7, Fig. S10

### The ARRIVE guidelines 2.0: author checklist

#### The ARRIVE Essential 10

These items are the basic minimum to include in a manuscript. Without this information, readers and reviewers cannot assess the reliability of the findings.

| Item | | Recommendation | Section/line number, or reason for not reporting |
|---|---|---|---|
| **Study design** | 1 | For each experiment, provide brief details of study design including: | Fig. 1 |
| | | a. The groups being compared, including control groups. If no control group has been used, the rationale should be stated. | |
| | | b. The experimental unit (e.g. a single animal, litter, or cage of animals). | Unit = 1 rat |
| **Sample size** | 2 | a. Specify the exact number of experimental units allocated to each group, and the total number in each experiment. Also indicate the total number of animals used. | Fig. 1 & Fig. S10 |
| | | b. Explain how the sample size was decided. Provide details of any *a priori* sample size calculation, if done. | Methods |
| **Inclusion and exclusion criteria** | 3 | a. Describe any criteria used for including and excluding animals (or experimental units) during the experiment, and data points during the analysis. Specify if these criteria were established *a priori*. If no criteria were set, state this explicitly. | Methods |
| | | b. For each experimental group, report any animals, experimental units or data points not included in the analysis and explain why. If there were no exclusions, state so. | Table 4; Fig. S10 |
| | | c. For each analysis, report the exact value of *n* in each experimental group. | Results; Fig. S10 |
| **Randomisation** | 4 | a. State whether randomisation was used to allocate experimental units to control and treatment groups. If done, provide the method used to generate the randomisation sequence. | Yes, see Methods |
| | | b. Describe the strategy used to minimise potential confounders such as the order of treatments and measurements, or animal/cage location. If confounders were not controlled, state this explicitly. | Not controlled |
| **Blinding** | 5 | Describe who was aware of the group allocation at the different stages of the experiment (during the allocation, the conduct of the experiment, the outcome assessment, and the data analysis). | Only technicians |
| **Outcome measures** | 6 | a. Clearly define all outcome measures assessed (e.g. cell death, molecular markers, or behavioural changes). | Methods |
| | | b. For hypothesis-testing studies, specify the primary outcome measure, i.e. the outcome measure that was used to determine the sample size. | Methods |
| **Statistical methods** | 7 | a. Provide details of the statistical methods used for each analysis, including software used. | Methods |
| | | b. Describe any methods used to assess whether the data met the assumptions of the statistical approach, and what was done if the assumptions were not met. | Methods |
| **Experimental animals** | 8 | a. Provide species-appropriate details of the animals used, including species, strain and substrain, sex, age or developmental stage, and, if relevant, weight. | Methods; Table 2 |
| | | b. Provide further relevant information on the provenance of animals, health/immune status, genetic modification status, genotype, and any previous procedures. | Methods |
| **Experimental procedures** | 9 | For each experimental group, including controls, describe the procedures in enough detail to allow others to replicate them, including: | Methods |
| | | a. What was done, how it was done and what was used. | |
| | | b. When and how often. | Methods |
| | | c. Where (including detail of any acclimatisation periods). | Methods |
| | | d. Why (provide rationale for procedures). | Methods |
| **Results** | 10 | For each experiment conducted, including independent replications, report: | Results; Tables 5-7 |
| | | a. Summary/descriptive statistics for each experimental group, with a measure of variability where applicable (e.g. mean and SD, or median and range). | |
| | | b. If applicable, the effect size with a confidence interval. | Results; Tables 5-7 |

Fig. S1, 1 of 2.

#### The Recommended Set

These items complement the Essential 10 and add important context to the study. Reporting the items in both sets represents best practice.

| Item | | Recommendation | Section/line number, or reason for not reporting |
|---|---|---|---|
| **Abstract** | 11 | Provide an accurate summary of the research objectives, animal species, strain and sex, key methods, principal findings, and study conclusions. | Abstract |
| **Background** | 12 | a. Include sufficient scientific background to understand the rationale and context for the study, and explain the experimental approach. | Introduction |
| | | b. Explain how the animal species and model used address the scientific objectives and, where appropriate, the relevance to human biology. | Introduction |
| **Objectives** | 13 | Clearly describe the research question, research objectives and, where appropriate, specific hypotheses being tested. | Introduction |
| **Ethical statement** | 14 | Provide the name of the ethical review committee or equivalent that has approved the use of animals in this study, and any relevant licence or protocol numbers (if applicable). If ethical approval was not sought or granted, provide a justification. | Methods |
| **Housing and husbandry** | 15 | Provide details of housing and husbandry conditions, including any environmental enrichment. | Methods |
| **Animal care and monitoring** | 16 | a. Describe any interventions or steps taken in the experimental protocols to reduce pain, suffering and distress. | Methods |
| | | b. Report any expected or unexpected adverse events. | Results |
| | | c. Describe the humane endpoints established for the study, the signs that were monitored and the frequency of monitoring. If the study did not have humane endpoints, state this. | Methods |
| **Interpretation/ scientific implications** | 17 | a. Interpret the results, taking into account the study objectives and hypotheses, current theory and other relevant studies in the literature. | Discussion |
| | | b. Comment on the study limitations including potential sources of bias, limitations of the animal model, and imprecision associated with the results. | Discussion |
| **Generalisability/ translation** | 18 | Comment on whether, and how, the findings of this study are likely to generalise to other species or experimental conditions, including any relevance to human biology (where appropriate). | Discussion |
| **Protocol registration** | 19 | Provide a statement indicating whether a protocol (including the research question, key design features, and analysis plan) was prepared before the study, and if and where this protocol was registered. | Yes; not registered |
| **Data access** | 20 | Provide a statement describing if and where study data are available. | Supplemental Info |
| **Declaration of interests** | 21 | a. Declare any potential conflicts of interest, including financial and non-financial. If none exist, this should be stated. | None |
| | | b. List all funding sources (including grant identifier) and the role of the funder(s) in the design, analysis and reporting of the study. | Acknowlegements |

Fig. S1, 2 of 2

*Product Data*

# D12451

#### Description

Rodent Diet with 45% kcal% fat.

#### Used in Research

Obesity
Diabetes

#### Packaging

Product is packed in 12.5 kg box. Each box is identified with the product name, description, lot number and expiration date.

#### Lead Time

IN-STOCK. Ready for next day shipment.

#### Gamma-Irradiation

Yes. Add 10 days to delivery time.

#### Form

Pellet, Powder, Liquid

#### Shelf Life

Most diets require storage in a cool dry environment. Stored correctly they should last 3-6 months.

#### Control Diets

D12450B

### Formula

| Product # | D12451 | |
|---|---|---|
| | gm% | kcal% |
| Protein | 24 | 20 |
| Carbohydrate | 41 | 35 |
| Fat | 24 | 45 |
| Total | | 100 |
| kcal/gm | 4.73 | |

| Ingredient | gm | kcal |
|---|---|---|
| Casein, 80 Mesh | 200 | 800 |
| L-Cystine | 3 | 12 |
| Corn Starch | 72.8 | 291 |
| Maltodextrin 10 | 100 | 400 |
| Sucrose | 172.8 | 691 |
| Cellulose, BW200 | 50 | 0 |
| Soybean Oil | 25 | 225 |
| Lard* | 177.5 | 1598 |
| Mineral Mix S10026 | 10 | 0 |
| DiCalcium Phosphate | 13 | 0 |
| Calcium Carbonate | 5.5 | 0 |
| Potassium Citrate, 1 H2O | 16.5 | 0 |
| Vitamin Mix V10001 | 10 | 40 |
| Choline Bitartrate | 2 | 0 |
| FD&C Red Dye #40 | 0.05 | 0 |
| **Total** | **858.15** | **4057** |

Formulated by E. A. Ulman, Ph.D., Research Diets, Inc., 8/26/98 and 3/11/99.

*Typical analysis of cholesterol in lard = 0.95 mg/gram.
Cholesterol (mg)/4057 kcal = 168.6
Cholesterol (mg)/kg = 196.5

RESEARCH DIETS INC.
www.ResearchDiets.com

**Research Diets, Inc.**
20 Jules Lane
New Brunswick, NJ 08901


Copyright © 2006 Research Diets, Inc. All rights reserved. D12451

Fig. S2. High-fat diet.

### D12450B, D12451, D03091001 and D03091001G

#### Rodent Diet With 10 or 45 kcal% Fat and Modification With 40 kcal% Protein

| **Product #** | **D12450B** | | **D12451** | | **D03091001** | | **D03091001G** | |
|---|---|---|---|---|---|---|---|---|
| % | gm | *kcal* | gm | *kcal* | gm | *kcal* | gm | *kcal* |
| Protein | 19.2 | *20* | 23.7 | *20* | 38.5 | *40* | 38.5 | *40* |
| Carbohydrate | 67.3 | *70* | 41.4 | *35* | 48.1 | *50* | 48.1 | *50* |
| Fat | 4.3 | *10* | 23.6 | *45* | 4.3 | *10* | 4.3 | *10* |
| Total | | *100* | | *100* | | *100* | | *100* |
| kcal/gm | 3.85 | | 4.73 | | 3.85 | | 3.85 | |
| **Ingredient** | **gm** | ***kcal*** | **gm** | ***kcal*** | **gm** | ***kcal*** | **gm** | ***kcal*** |
| Casein, 80 Mesh | 200 | *800* | 200 | *800* | 400 | *1600* | 400 | *1600* |
| L-Cystine | 3 | *12* | 3 | *12* | 6 | *24* | 6 | *24* |
| Corn Starch | 315 | *1260* | 72.8 | *291* | 213.5 | *854* | 213.5 | *854* |
| Maltodextrin 10 | 35 | *140* | 100 | *400* | 35 | *140* | 35 | *140* |
| Sucrose | 350 | *1400* | 172.8 | *691* | 248.5 | *994* | 248.5 | *994* |
| Cellulose, BW200 | 50 | *0* | 50 | *0* | 50 | *0* | 50 | *0* |
| Soybean Oil | 25 | *225* | 25 | *225* | 25 | *225* | 25 | *225* |
| Lard | 20 | *180* | 177.5 | *1598* | 20 | *180* | 20 | *180* |
| Mineral Mix S10026 | 10 | *0* | 10 | *0* | 10 | *0* | 10 | *0* |
| DiCalcium Phosphate | 13 | *0* | 13 | *0* | 13 | *0* | 13 | *0* |
| Calcium Carbonate | 5.5 | *0* | 5.5 | *0* | 5.5 | *0* | 5.5 | *0* |
| Potassium Citrate, 1 H2O | 16.5 | *0* | 16.5 | *0* | 16.5 | *0* | 16.5 | *0* |
| Vitamin Mix V10001 | 10 | *40* | 10 | *40* | 10 | *40* | 10 | *40* |
| Choline Bitartrate | 2 | *0* | 2 | *0* | 2 | *0* | 2 | *0* |
| FD&C Yellow Dye #5 | 0.05 | *0* | 0 | *0* | 0 | *0* | 0.025 | *0* |
| FD&C Red Dye #40 | 0 | *0* | 0.05 | *0* | 0 | *0* | 0 | *0* |
| FD&C Blue Dye #1 | 0 | *0* | 0 | *0* | 0 | *0* | 0.025 | *0* |
| **Total** | **1055.05** | ***4057*** | **858.15** | ***4057*** | **1055** | ***4057*** | **1055.05** | ***4057*** |

Formulated by Research Diets, Inc.

Fig. S3. Low-fat diet (D12450B).

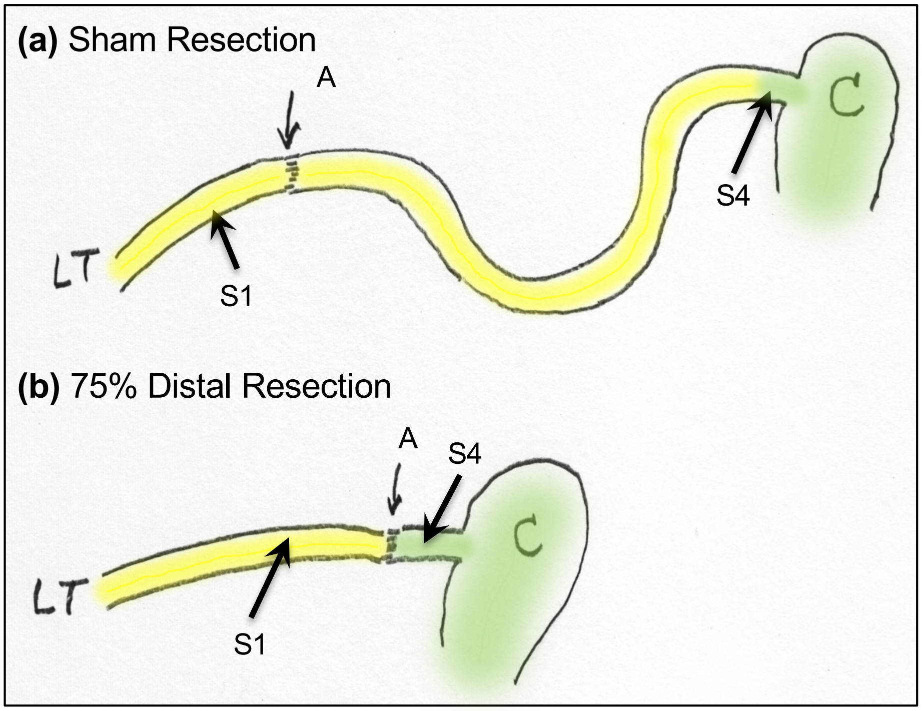

**Fig. S4. Resection Scheme.**

**(a)** Sham resection. LT = Ligament of Trietz; A = transection and anastomotic site; C = cecum; S1 = proximal jejunum; S4 = distal ileum. The local of site A is the junction of the proximal 25% of the small intestine and the distal 75% of the small intestine (distances determined by measuring intestine with a cotton umbilical tape).

**(b)** 75% Distal (massive) resection. The proximal transection site is "A" from panel (a); the distal transection site is in the distal ileum, one cm from the cecum (i.e., the junction of the yellow and green-shaded segments). The intervening segment (consisting of ~half of the jejunum and nearly all of the ileum, representing 75% of the entire small bowel length) is excised, and the proximal jejunum is anastomosed to the distal ileum.

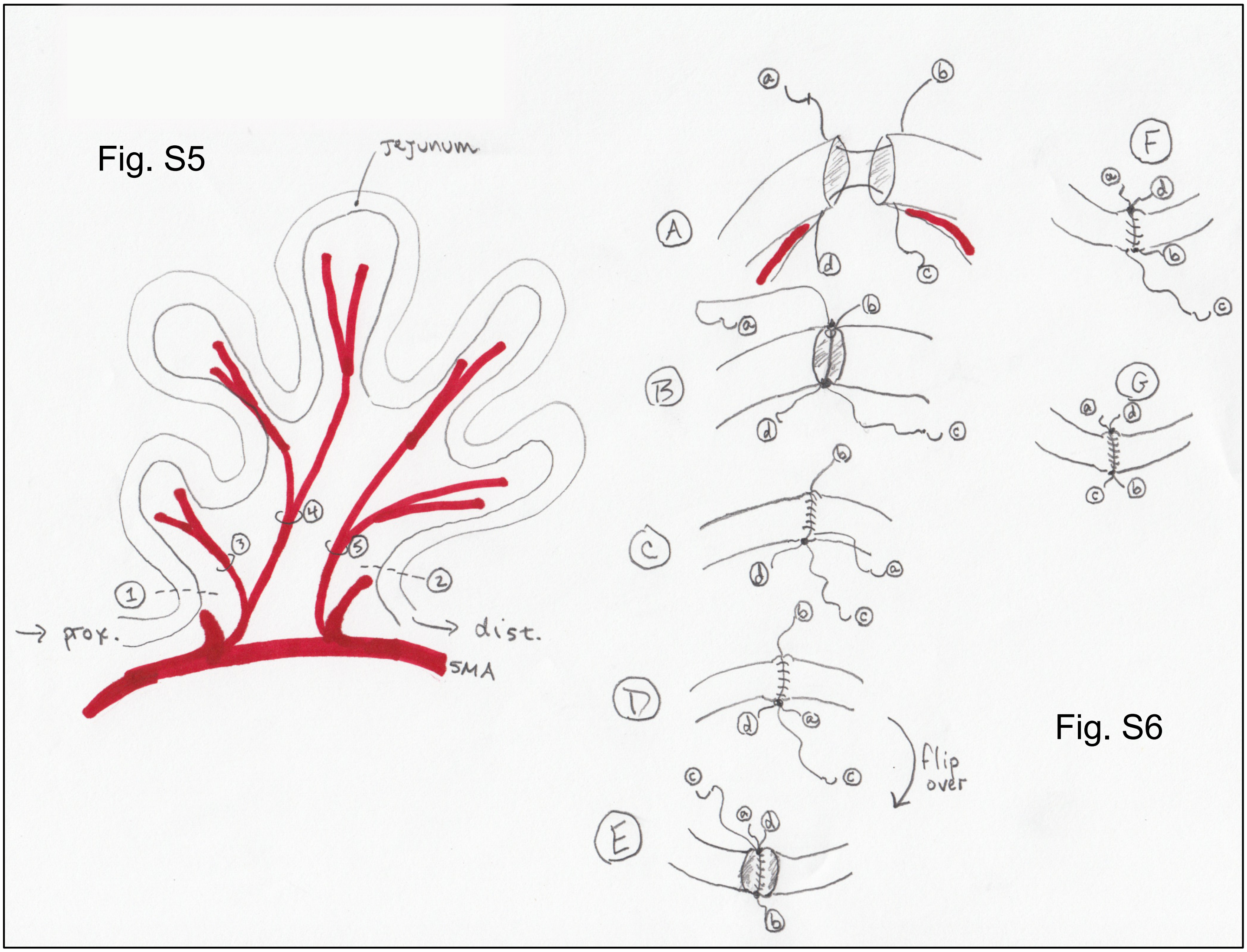

**Fig. S5. Resection technique**. Proximal & distal points of transection (1 & 2) can be slightly adjusted to be adjacent to feeding vessels, so that transected intestinal ends will be well-vascularized. Ligation of mesenteric vessels (3-5) is done somewhat away from mesenteric root in order to minimize risk of injury to superior mesenteric artery (SMA) injury. Other general points:

- Avoid stretching of mesentery
- Keep all intestine (other than that being resected and the ends being sewn together) inside the abdomen during procedure
- Keep surface of retained bowel wet with saline
- Use sponge under the anastomosis during suturing to catch any intestinal spillage

**Fig S6. Anastomotic technique**. Perform enteroenterostomy with 6-0 or silk, double armed, cut in half. (A) Line up two ends of intestine as shown; using the two suture halves, place corner stitches at mesenteric and anti-mesenteric borders; leave tails (b & d) long enough for tying. Make sure that mesenteric feeding vessels (in red) reach the end of each intestinal limb. (B) Tie corner stitches. (C) Run arm “a” to create anterior anastomotic suture line, using full thickness bites, outside-in to inside-out (i.e. sewing on the bowel exterior). (D) Tie arm “a” to tail “d”. (E) Flip intestines over to create posterior suture line. (F) Run arm “c” to create posterior anastomotic suture line, similar to panel C. (G) Tie arm “c” to tail “b” to complete anastomosis.

Formulated by:
Research Diets, Inc.

### Rodent Diets with 45 kcal% Fat or 25 kcal% Fat with 17 kcal% Sucrose

| Product # | D12451 | | D14041301 | |
|---|---|---|---|---|
| % | gm | *kcal* | gm | *kcal* |
| Protein | 24 | *20* | 21 | *20* |
| Carbohydrate | 41 | *35* | 58 | *55* |
| Fat | 24 | *45* | 12 | *25* |
| Total | | *100* | | *100* |
| kcal/gm | 4.7 | | 4.2 | |
| **Ingredient** | **gm** | ***kcal*** | **gm** | ***kcal*** |
| Casein | 200 | *800* | 200 | *800* |
| L-Cystine | 3 | *12* | 3 | *12* |
| Corn Starch | 72.8 | *291* | 276.5 | *1106* |
| Maltodextrin 10 | 100 | *400* | 100 | *400* |
| Sucrose | 172.8 | *691* | 172.8 | *691* |
| Cellulose, BW200 | 50 | *0* | 50 | *0* |
| Soybean Oil | 25 | *225* | 25 | *225* |
| Lard | 177.5 | *1598* | 87 | *783* |
| Mineral Mix S10026 | 10 | *0* | 10 | *0* |
| DiCalcium Phosphate | 13 | *0* | 13 | *0* |
| Calcium Carbonate | 5.5 | *0* | 5.5 | *0* |
| Potassium Citrate, 1 H2O | 16.5 | *0* | 16.5 | *0* |
| Vitamin Mix V10001 | 10 | *40* | 10 | *40* |
| Choline Bitartrate | 2 | *0* | 2 | *0* |
| FD&C Yellow Dye #5 | 0 | *0* | 0.025 | *0* |
| FD&C Red Dye #40 | 0.05 | *0* | 0 | *0* |
| FD&C Blue Dye #1 | 0 | *0* | 0.025 | *0* |
| **Total** | **858.15** | ***4057*** | **971.35** | ***4057*** |

Research Diets, Inc.
20 Jules Lane
New Brunswick, NJ 08901 USA


Fig. S7. Intermediate-fat diet (D14041301).

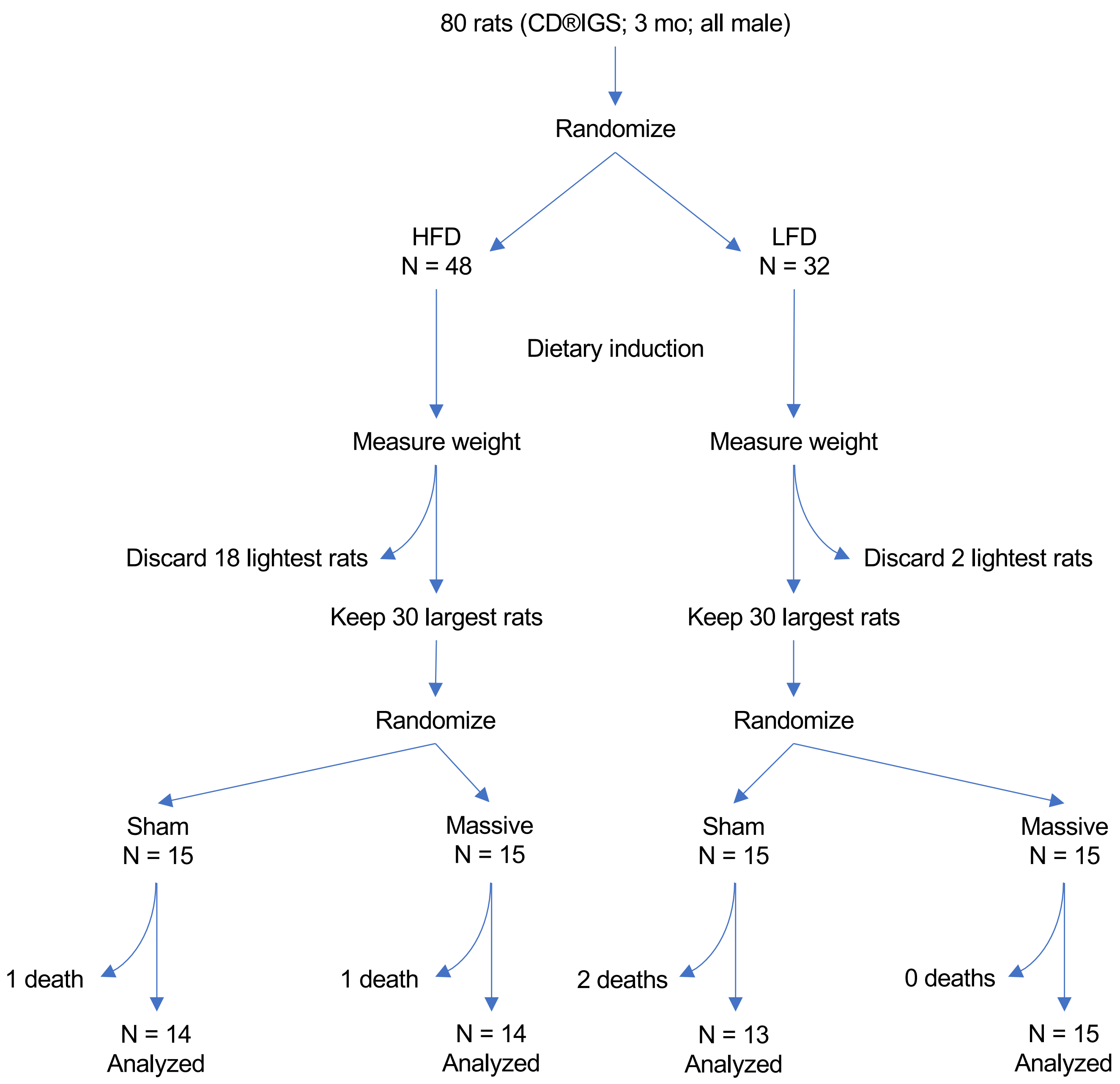

**Fig. S10**. Actual fate of all 80 rats entered into the study. HFD = high fat diet; LFD = low fat diet; sham = sham resection; Massive = 75% distal small intestinal resection.
